# Identification of a Major Locus for Flowering Pattern Sheds Light on Plant Architecture Diversification in Cultivated Peanut

**DOI:** 10.1101/2021.03.18.435916

**Authors:** Srinivas Kunta, Ye Chu, Yael Levy, Arye Harel, Shahal Abbo, Peggy Ozias-Akins, Ran Hovav

## Abstract

Flowering pattern is a major taxonomic characteristic differentiating the two main subspecies of cultivated peanut (*Arachis hypogaea* L.). subsp. *fastigiata* possessing flowers on the mainstem (MSF) and a sequential flowering pattern, whereas subsp. *hypogaea* lacks flowers on the mainstem and exhibits an alternate flowering pattern. This character is considered the main contributor to plant architecture and the adaptability of each subgroup to specific growing conditions. Evidence indicates that flowering pattern differentiation occurred during the several thousand years of domestication and diversification in South America. However, the exact genetic mechanism that controls flowering pattern and the molecular changes that led to its historical diversification in peanut are unknown. We investigated the genetics of the flowering pattern in a recombinant inbred population of 259 lines (RILs), derivatives of an *A. hypogaea* and *A. fastigiata* cross. RILs segregated 1:1 in both the sequential/alternative and the MSF-plus/MSF-minus traits, indicating a single gene effect. Using the Axiom_Arachis2 SNP-array, MSF was mapped to a 1.7 Mbp segment on chromosome B02 of the cultivated *A. hypogaea*. Significant haplotype conservation was found for this locus in the USA peanut mini core collection, suggesting a possible selection upon *hypogaea*/*fastigiata* speciation. Furthermore, a candidate *Terminal Flowering 1*-*like* (*AhTFL1*) gene was identified within the MSF region, in which a 1492 bp deletion occurred in the *fastigiata* line that leads to a truncated protein product. Remapping MSF in the RIL population with the *AhTFL1* deletion as a marker increased the LOD score from 53.3 to 158.8 with no recombination. The same deletion was also found to co-segregate with the phenotype in two EMS-mutagenized M2 families, suggesting a hotspot for large mutational deletion or gene conversion that may play a role in evolution. BLASTX analysis showed that the most similar homologous gene for *TFL1-like* in soybean is *Det1*, which previously was shown to control shoot determination. Sequence analysis of the *TFL-1* in a series of domesticated lines showed that *TFL1* was subjected to gain/loss events of the deletion, partly explaining the evolution of MSF in Arachis. Altogether, these results support the role of *AhTFL-1* in peanut speciation during domestication and modern cultivation.

## Introduction

Cultivated peanut (*Arachis hypogaea* L.) or groundnut (*A. hypogaea* L.) is among the most important oil and food legume crops. Peanut is produced in over 100 countries, mainly in tropical and subtropical zones (1). It is consumed worldwide and therefore also plays a substantial role in global trade. As a nitrogen-fixing legume, peanut is a vital rotation crop that restores nitrogen to the soil. Cultivated peanut is an allotetraploid species (AABB) domesticated in central South America. It originated from the hybridization of two wild progenitor diploid species: *A. duranensis* (AA) and *A. ipaënsis* (BB), followed by genome duplication (2, 3). The cultivated peanut genome size is about 2.7 Gb, an approximate sum of the two diploid A- and B-genome wild progenitors (4). The polyploidization event was relatively recent, ~9-10 thousand years ago at most (3), which speciated the cultivated peanut from its wild diploid relatives.

Few distinct botanical subspecies were established during several thousand years of peanut domestication and diversification in South America. The two main subspecies, which are currently grown worldwide, are the *A. hypogaea* subsp. *fastigiata* (that includes the ‘Spanish’ and ‘Valencia’ market-types), and the *A. hypogaea* subsp. *hypogaea* (that includes the ‘Virginia’ and ‘Runner’ market-types). Archaeological records indicate that these two subspecies originated in South American regions corresponding with today’s Chile, Argentina, Ecuador, Paraguay, Bolivia, and Brazil (5). General notion suggests that *fastigiata* was evolved from *hypogaea*, but this has occurred almost immediately upon the later was developed from a common ancestor. Other botanical types include *peruviana, aequatoriana*, and *hirsuta* that were developed in the northern part of South America (Peru, Bolivia, and Ecuador) under the respective subspecies (6) but are not being cultivated on a large scale.

Subspecies *fastigiata* and *hypogaea* are distinct by marketing, agronomic and morphological features. From a marketing perspective, pod characteristics such as pod size, seed size, and seed coat color are the phenotypes used to distinguish *fastigiata* from *hypogaea* (7). From an agronomic perspective, the main difference between the *fastigiata* and *hypogaea* is the growing period. Varieties from the *fastigiata* subspecies have short-duration growing time (from 90-120 days), while the *hypogaea*-related varieties are usually late-maturing (130-170 days) (8). Therefore, *fastigiata* is being grown in low input agricultural systems in regions with a short rainy season or two seasons per year, such as in Africa, South-East Asia, and Central America. On the other hand, *hypogaea* cultivars are usually high-yielding and are being grown in regions with a long summer, frequently with supplemental irrigation, like in the USA and the Middle East. In 2019, *fastigiata* peanuts constituted ~90% of the global peanut production (~42.5 Million tons) (1).

The major taxonomic feature distinguishing the two subspecies is the type of growth habit. First, the lateral shoots of the *fastigiata* types are erect (which is the meaning of the Latin name *fastigiata*), while *hypogaea* types have spreading or bunch laterals. Yet, the most common morphological feature distinguishing *fastigiata* from hypogaea is the flowering pattern. Subspecies *fastigiata* possesses flowers on the mainstem (MSF) and a sequential branching pattern, whereas subspecies *hypogaea* lacks flowers on the mainstem and exhibits an alternate branching pattern (**Fig. 1**). The flowering pattern is the factor that controls plant shoot determination, and therefore it is the main effect for the difference in time-to-maturity between *fastigiata* and *hypogaea* (8). It is almost certain that the sequential flowering formation’s historical event was an essential prerequisite for the domination of *fastigiata* in areas with a short rainy season or high drought stress level. As a result, the flowering pattern has become a primary descriptive trait for peanut (9) and an integral part of variety characterization.

**Figure 1.**
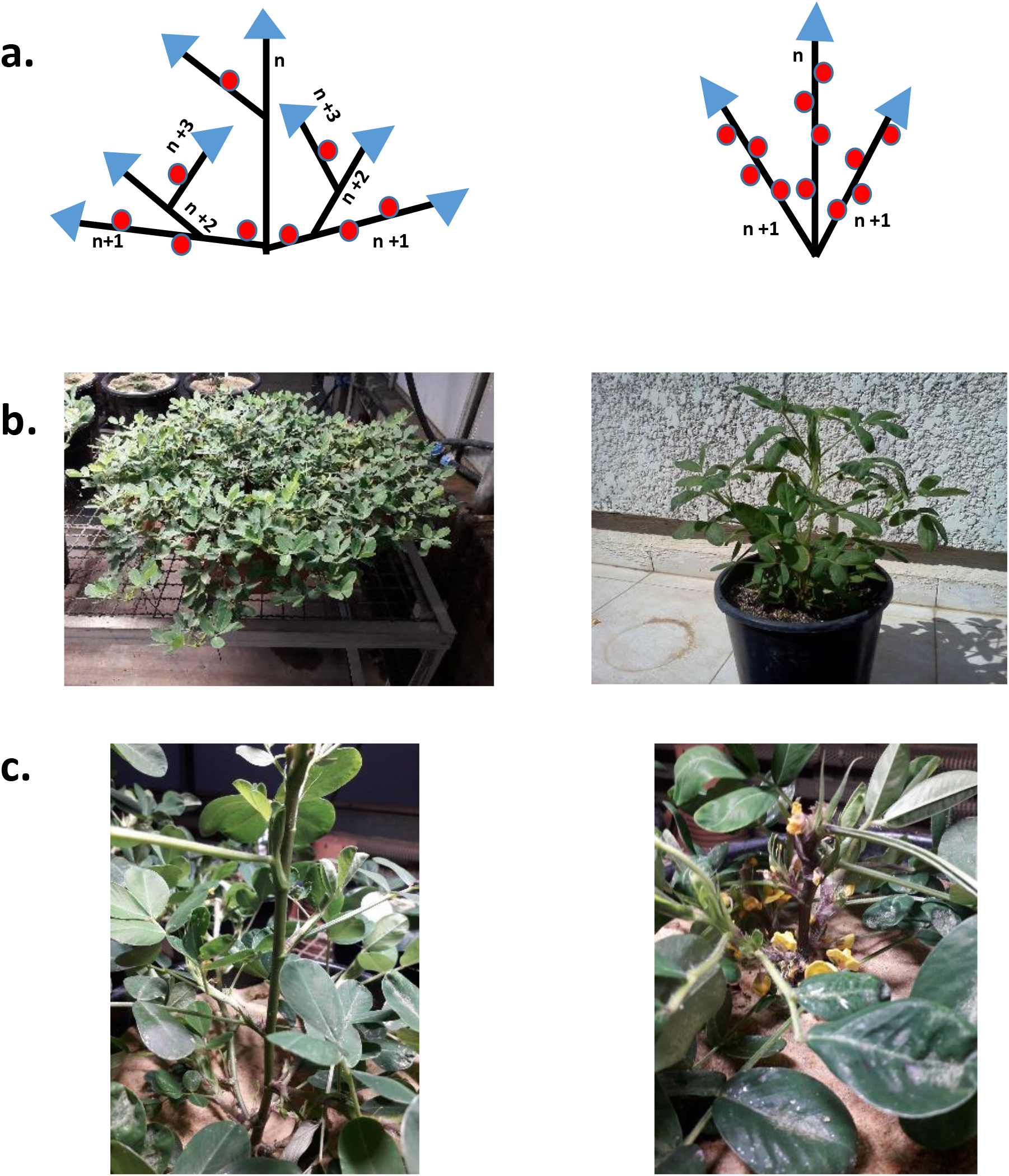
Comparing the growth habits between Hanoch (left) and IGC99 (right), the two parental genotypes represent *A. hypogaea* subsp. *hypogaea* and *A. hypogaea* subsp. *fastigiata*, respectively. a. A schematic diagram of the two flowering patterns (left = Hanoch; right = IGC99). Triangles indicate the growth axis from meristems; Indeterminate meristems are colored blue; balls represent determinate floral buds. **b**. Real-life example photos of the two genotypes. **c**. The absence and the presence of flowers on the main stem of Hanoch (left) and IGC99 (right), respectively.

Despite its descriptive and agronomic importance, currently, not much is known about the genetic control of the flowering pattern in peanut. Also, the evolutionary events that led to the occurrence of the *fastigiata* sequential flowering pattern are mainly unknown. Previous studies have reported that the MSF trait in the cultivated peanut is controlled by two sets of duplicate loci interacting with epistasis (10, 11). Chopra et al. (12) have identified a major QTL effect (44%) on LG6 for the presence of flowers on a mainstem in a cross-system that involves two A-genome wild species (*A. duranensis* and *A. cardenasii*). They also identified two additional putative QTLs on LG8 explaining 10% of phenotypic variance each. The *A. cardenasii* was the donor parent of the MSF trait in that study, while *A. ipaensis* is considered the A-genome diploid progenitor of the cultivated peanut.

The current study aimed to identify the genetic factor that controls flowering pattern in cultivated peanut. A Recombinant Inbred Line (RIL) population, derived from a *fastigiata* × *hypogaea* cross, was analyzed to determine the mode of inheritance for the flowering pattern trait and to identify genomic loci that control it. Using a powerful SNP-array technology, we succeeded in identifying a major locus that controls the sequential flowering pattern and MSF in our genetic background. An association study of this locus in the peanut mini core collection and candidate gene analyses were performed to obtain new insight into the molecular evolution of this trait in *A. hypogaea*.

## Materials and Methods

### Plant material and growing conditions

A recombinant inbred line (RIL) population was advanced from a cross between cv. ‘Hanoch’ and cv. ‘Congo-Red’ (IGC99). Hanoch is a Virginia-type cultivar (*A. hypogaea* subsp. *hypogaea* var. *hypogaea*), the leading Israeli in-shell peanut cultivar for three decades. It is a late-maturing spreading cultivar with large pods. Hanoch has an alternate flowering pattern with no visible flowers on the main stem (MSF-minus) (**Fig. 1a,c**). IGC99 (Israeli Groundnut Collection No. 99) is an old Valencia-type (*A. hypogaea* subsp. *fastigiata* var. *fastigiata*) cultivar that was commercially grown in Israel during 1970-1980. It possesses erect lateral shoots, sequential flowering and flowers on the main stem (MSF-plus) (**Fig. 1a,c**).

A total of 259 RILs were planted in April 2020 within a commercial plot at Nir Itzhak, Western Negev, Israel. A randomized complete block experimental design with three replications was implemented. Each ‘plot’ (RIL X block) consisted of two rows on a bed, 4 m in length, rows spaced 90 cm apart and seeding rates of 10 seeds/m^2^ (total of 20 plants/plot). Parental lines were grown as control plots with nine replications. Fields were maintained under full-irrigation conditions.

A previously developed Hanoch-based ethyl methane sulfonate (EMS) - mutagenized population (13) was used to analyze the phenotype and the genotype of MSF mutants. Four M_2_ families previously found to have at least 1 out of 14 EMS mutants were analyzed. From each family, ~15 plants were grown in 5L pots in a greenhouse DNA was extracted from all plants.

### Phenotyping the flowering and branching patterns in the RIL population

Phenotyping of the flowering pattern and the branching habit traits was done at 70 days post-planting. All 20 plants within a plot were phenotyped for the presence of flowering on the main stem (MSF-plus vs. MSF-minus). If a RIL was still segregating for MSF, it was excluded from the general analysis. Still, the pattern of MSF segregation within these segregating RILs was scored on a single plant basis.

In the same manner, the flowering pattern, i.e., sequential vs. alternative, was scored. In alternative flowering, the lateral shoots (n+1) are flowering, and the secondary lateral shoots (n+2) are formed without flowers. The next level of lateral shoots (n+3) bear flowers and so on (**Fig 1a**). The plant, therefore, exhibits many lateral branches (**Fig. 1b**). In the sequential flowering pattern, there are flowers on the lateral n+1 shoots, but the n+1 shoots are not formed, and the axillary bud is terminated into a flower. The plant exhibits only 4-5 branches (**Fig. 1b**). Therefore, to facilitate rapid phenotyping, the flowering pattern was scored as branching rate (BR), wherein the alternative was defined as “Many laterals” and the sequential as “up to 5 laterals”. To inspect the association between flowering pattern and branching habit (BH), the growing angle of the lateral shoots was recorded by defining lines as “spreading”, “bunch”, or “erect” according to the US peanut descriptor guidelines (9). The chi-square test of the JMP^®^ Pro 15 software (SAS Institute Inc., Cary, NC, 1989-2019) was used to analyze the traits’ segregation pattern.

### Genotyping and genetic map construction

Genomic DNA was extracted from young leaflets of each RIL and the two parents using DNeasy® Plant Mini Kit (Qiagen; Hilden, Germany). DNA quantification was performed with Qubit (Invitrogen; CA, USA). The samples were diluted to 40 ng/μL according to protocol guidelines and genotyped using the Affymetrix Axiom_Arachis2 SNP-array comprising 47,837 SNPs (14, 15).

Genotyping data were analyzed by the Axiom analysis suite Software 3.1, as previously described (16) with some modifications. The polymorphic SNPs (AA or BB and AB) were retained with 35–65% call-rate frequencies among the RILs. Markers with more than 10% missing data and more than 20% heterozygote calls (in AA/BB parental SNPs) were removed. The genetic linkage map was constructed using Joinmap v4.1 (17) with a minimum LOD of 3.0 and the Kosambi mapping function. The graphical representation of the linkage maps was generated through MapChart v2.3 (18). Confirmation of the loci positions was done as previously described (16) with few modifications (BLASTN (e value <1×10-18) and mismatch of less than 2). Generated linkage groups (LG) were assigned to the pseudo-molecules of the tetraploid *A. hypogaea* cv. Tifrunner (19). To evaluate the quality of the genetic map, a colinearity analysis was performed using the genetic distances (cM) versus the physical positions (Mbp) plot.

### Trait Mapping

Mapping of MSF, BR and the BH was performed on the 259 RILs using MapQTL v6 (20) by converting each qualitative phenotype into a number. For example, MSF-plus =1 vs. MSF-minus = 2. Loci were detected by interval mapping using significance threshold levels derived from permutation tests. SNP markers flanking the QTL were used to obtain the physical position from the *A. hypogaea* genome (peanutbase.org).

### Determination of haplotype conservation in the US mini core collection

A haplotype conservation analysis was performed to determine the level of conservation flanking MSF in the US peanut mini core collection. Phenotyping of the collection for MSF was done by evaluating the presence or absence of flowers on individual plants grown in the greenhouse at 106 days after planting. Two tightly linked markers (AX-147227990, AX-176805840) that demonstrated polymorphic pattern within the mini core collection were chosen. Significant linkage disequilibrium was found within this MSF region. Out of 103 accessions, only eleven accessions demonstrated heterozygosity and/or recombination among the two markers. The remaining 92 accessions were included in the haplotype analysis, among which 44 and 48 exhibited Hanoch and IGC99 haplotypes, respectively. For haplotype conservation analysis, each allele’s frequencies were calculated in each group, and a chi-square analysis was performed between the frequencies of the two groups. The H_0_ is that no significant differences should be detected between the allele frequencies. The H_0_ is rejected at prob(ChiSq)<0.05.

### Sequencing TFL-1 like gene and its deletion

To clone the deletion of the *TFL-1* like gene, two sets of primers were designed based on the *TFL-1* sequence in the Tifrunner tetraploid genome (arahy.BBG51B.1; peanutbase.org). The first two primers, F-GGAAGGGAAAGAGATAGTGAGCT and R-GCATTATTCACTAATTGACATCACAAG (5 min initial denaturation at 95°C; 36 cycles of denaturation (30 s at 95°C), annealing (30 s at 62°C), and extension (90 s at 72°C); a final extension at 72°C for 5 min; Hy-Taq Ready Mix (2x)-hylabs^®^), were used to amplify across the complete deletion. The second set of primers was used to amplify the last exon of the *TFL-1* gene, which is absent in *fastigiata* and present in the *hypogaea* B-genome. The sequences of these primers are F-GCCACATTTGGTAGGTTTCATTG and R-CATTTCTTGGCTTGTAAACATGAC (5 min initial denaturation at 95°C; 36 cycles of denaturation (30 s at 95°C), annealing (30 s at 61°C), and extension (80 s at 72°C); a final extension at 72°C for 5 min; Hy-Taq Ready Mix (2x) - hylabs^®^).

## Results

### Trait phenotyping and genetic analyses

The RIL population, derived from a cross between Hanoch (*hypogaea*) X IGC99 (*fastigiata*), was scored for the presence of flowering on main steam (MSF), branching rate (BR) and branching habit (BH) (**Table S1**). Out of 269 lines, 259 had uniform MSF phenotype (i.e., MSF-plus or MSF-minus), while the rest 10 lines were still segregating for the trait. Out of the 259 uniform lines, 126 were MSF-plus, and 133 were MSF-minus. Chi-square test for a segregation ratio of 1:1 was not significant (Chi value = 0.189; p(chi) = 0.664). Moreover, analyzing the 10 still segregating lines (total of 196 plants), the MSF-minus: MSF-plus ratio was 141:55, which did not significantly deviate from 3:1 (Chi value = 0.471; P(chi) = 0.492). Therefore, we concluded that a single gene/locus controls the MSF trait in this genetic background.

The BR trait segregation in the RIL population completely matched MSF; i.e., lines with MSF-plus were “up to 5 laterals”, whereas MSF-minus scored as “Many laterals”. This result showed that the same genetic factor controls both MSF and BR. BH segregated within the RILs as follows: 139 were “spreading”, 81 “bunch”, 28 “erect”. No particular genetic model was found to explain the mode of this segregation. No correlation was found between the MSF/BR phenotype and the BH phenotype, suggesting no genetic linkage between MSF and BH in this RIL population.

### Construction of the genetic map

Genotyping of Hanoch x IGC99 RILs used the Axiom_ Arachis2 SNP-array. Out of ~47,000 SNPs on the chip, 8251 polymorphic SNPs were identified for the parental lines and the RILs. After filtering to remove the excess heterozygous calls (> 20%) and missing data (> 10%), 6870 SNPs were retained for the genetic map constructed with 259 RILs.

The constructed genetic map covered 2751.6 cM, containing 4725 markers allocated to 21 linkage groups (**Table S2**, **Fig. 2a**). Linkage groups were assigned to the respective pseudomolecules (chromosomes) of the sequenced *A. hypogaea* genome cv.Tifrunner (**Table 1**). If a LG had more than 51% of the SNPs representing a particular chromosome, this LG was assigned to that chromosome. Most linkage groups comprised markers from both homeologous A- and B- chromosomes, but the highest proportion was in LG A07 with 12% of the Arahy.12 chromosome. Most of the linkage groups had markers from other non-homeologous chromosomes, but altogether, the proportion did not exceed 5% in the respective linkage groups. The 21 linkage groups ranged in size from 15.09cM (B06) to 184.94 cM (B09). The average number of loci per linkage group was 225 reaching up to 462 loci in B09. The average distance between the neighboring markers was 0.65 cM, ranging from 0.43 cM in LGs A01 to 1.07 cM in A03_2 (**Table 1**).

**Figure 2.**
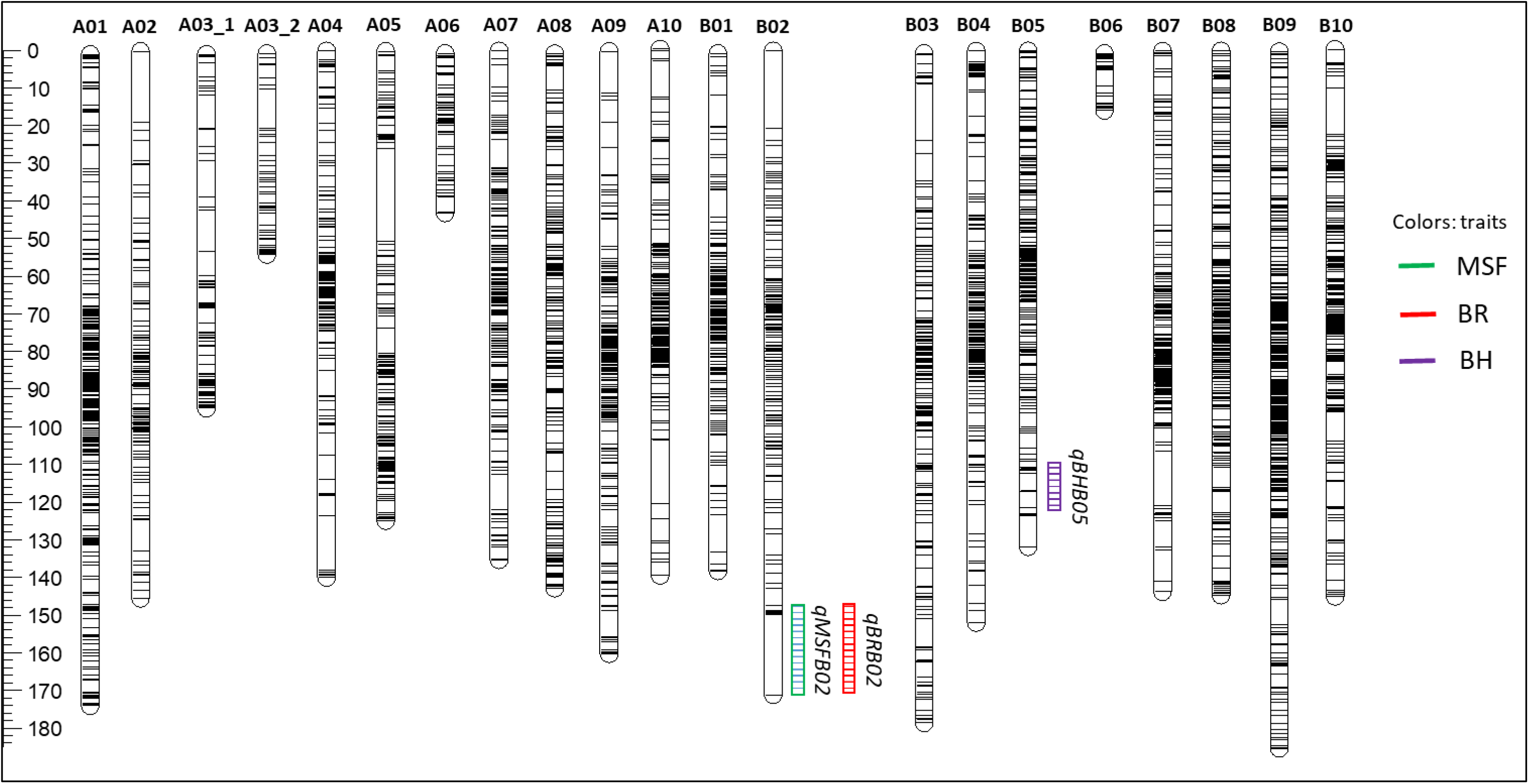
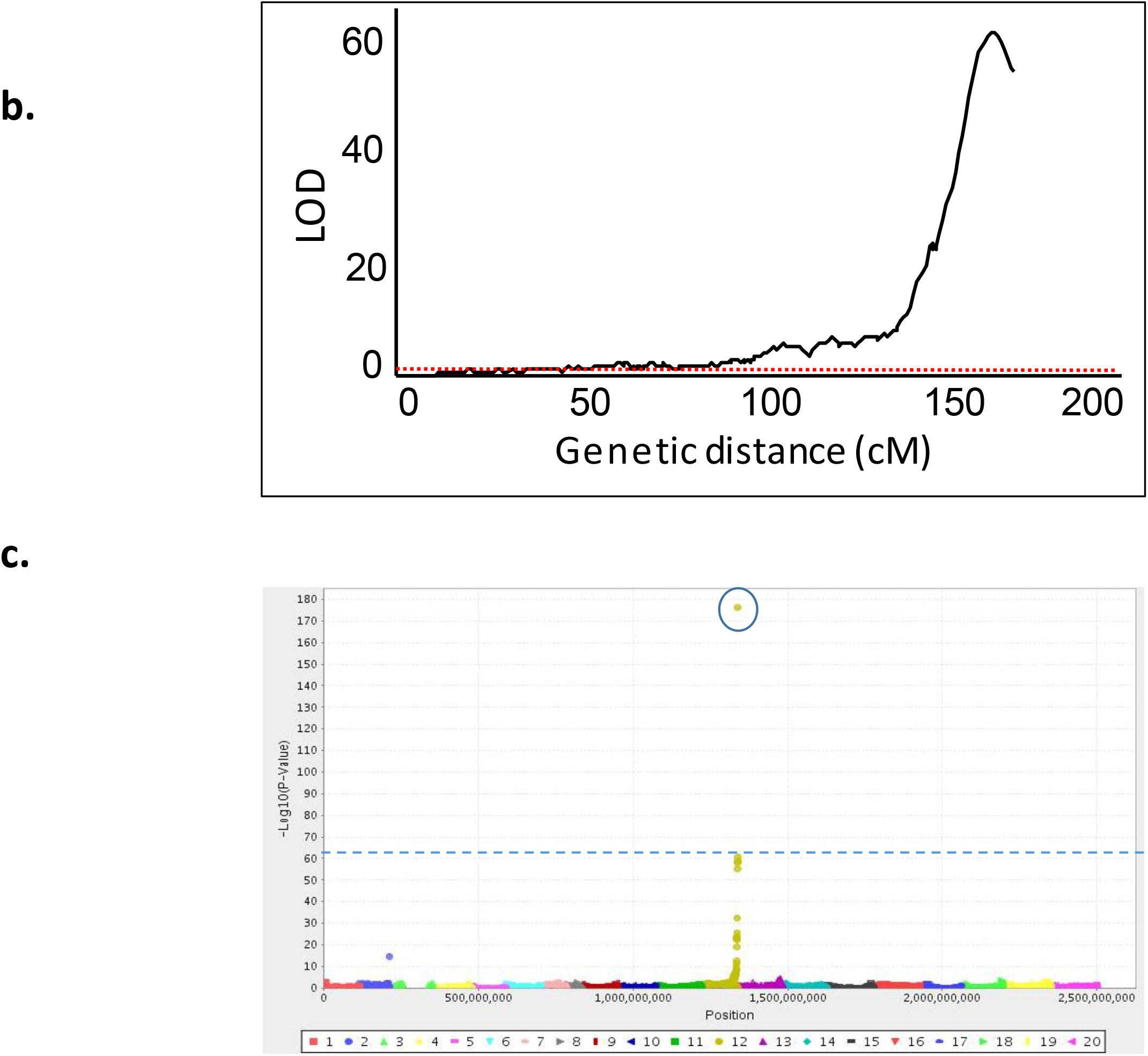
Mapping of MSF, BR and BH traits. **a**. A genetic map consisting of 4887 SNP markers, based on Hanoch X IGC99 RIL population. **b**. Genetic mapping of the MSF trait. **c**. Manhattan plot for the correlation between the allele of each of 6871 SNP markers and the MSF phenotype. **d**. Linkage groups with significant loci for MSF, BR and BH mapped with MapQTL 6.

**Table 1:** Features of the saturated genetic linkage map with 4887 SNP markers for Hanoch x IGC99 RIL population.

| Linkage group | Assigned Chromosome | No. of SNPs | Genetic length (cM) | Average loci interval | Physical Length (Mbp) | Average physical interval | Total length A. hypogaea genome | Coverage ratio | Recombination rate (cM/Mbp) |
| --- | --- | --- | --- | --- | --- | --- | --- | --- | --- |

|  |  |  |  | (cM) |  | (Mbp) | (Mbp) |  |  |
| --- | --- | --- | --- | --- | --- | --- | --- | --- | --- |
| A01 | Arahy.01 | 405 | 172.99 | 0.43 | 111.98 | 0.28 | 112.42 | 1.00 | 1.54 |
| A02 | Arahy.02 | 140 | 145.34 | 1.04 | 96.01 | 0.69 | 102.98 | 0.93 | 1.41 |
| A03_1 | Arahy.03 | 96 | 93.98 | 0.98 | 42.55 | 0.44 | 143.81 | 0.30 | 0.65 |
| A03_2 |  | 50 | 53.53 | 1.07 | 10.15 | 0.20 | 143.81 | 0.07 | 0.37 |
| A04 | Arahy.04 | 183 | 139.74 | 0.76 | 125.52 | 0.69 | 128.8 | 0.97 | 1.08 |
| A05 | Arahy.05 | 200 | 124.74 | 0.62 | 109.43 | 0.55 | 115.93 | 0.94 | 1.08 |
| A06 | Arahy.06 | 64 | 42.42 | 0.66 | 8.85 | 0.14 | 115.5 | 0.08 | 0.37 |
| A07 | Arahy.07 | 261 | 135.02 | 0.52 | 80.38 | 0.31 | 81.12 | 0.99 | 1.66 |
| A08 | Arahy.08 | 218 | 141.89 | 0.65 | 51.12 | 0.23 | 51.9 | 0.98 | 2.73 |
| A09 | Arahy.09 | 272 | 160.01 | 0.59 | 120.11 | 0.44 | 120.52 | 1.00 | 1.33 |
| A10 | Arahy.10 | 255 | 139.51 | 0.55 | 115.52 | 0.45 | 117.09 | 0.99 | 1.19 |
| B01 | Arahy.11 | 271 | 137.45 | 0.51 | 148.93 | 0.55 | 149.3 | 1.00 | 0.92 |
| B02 | Arahy.12 | 176 | 170.76 | 0.97 | 116.85 | 0.66 | 120.58 | 0.97 | 1.42 |
| B03 | Arahy.13 | 222 | 177.76 | 0.80 | 145.72 | 0.66 | 146.73 | 0.99 | 1.21 |
| B04 | Arahy.14 | 280 | 151.91 | 0.54 | 140.88 | 0.50 | 143.24 | 0.98 | 1.06 |
| B05 | Arahy.15 | 282 | 131.54 | 0.47 | 153.01 | 0.54 | 160.88 | 0.95 | 0.82 |
| B06 | Arahy.16 | 34 | 15.09 | 0.44 | 80.26 | 2.36 | 154.81 | 0.52 | 0.10 |
| B07 | Arahy.17 | 221 | 143.25 | 0.65 | 133.39 | 0.60 | 134.92 | 0.99 | 1.06 |
| B08 | Arahy.18 | 329 | 144.39 | 0.44 | 133.93 | 0.41 | 135.15 | 0.99 | 1.07 |
| B09 | Arahy.19 | 462 | 184.94 | 0.40 | 159.09 | 0.34 | 158.63 | 1.00 | 1.17 |
| B10 | Arahy.20 | 304 | 145.34 | 0.48 | 144.48 | 0.48 | 143.98 | 1.00 | 1.01 |
|  | Mean | 225 | 131.03 | 0.65 | 106.10 | 0.55 | 129.48 | 0.84 | 1.11 |
|  | Total | 4725 | 2751.60 | 13.56 | 2228.16 | 11.52 | 3107.48 | 17.65 | 23.25 |

The 4725 loci aligned to the *A. hypogaea* pseudomolecules spanned a total physical distance of 2228.16 Mbp and an average physical interval of 0.55 Mbp between loci (**Fig. S1**; **Table 1**). The percentage of pseudomolecules covered by the linkage map varied. Twelve groups covered more than 90%, and four groups were close to 100% of the pseudomolecule. The average recombination rate was 1.1 cM/Mbp. A08 and A01 had the maximum recombination rate, while B06 and A03_2 had the lowest recombination rates. The linkage map quality was assessed by analyzing the collinearity of SNPs to their physical positions (Mbp) in the A. hypogaea genome (**Fig. S1**). As expected, the saturation of the markers in the arms was higher than in the pericentromeric regions. Some rearrangements were exhibited in few linkage groups, such as apparent inversions in the middle of B09 (**Fig. S1**).

### Trait mapping

Mapping of the MSF, BR and BH is presented in **Fig 2**. *MSF* was mapped to a single genomic locus on LG B02, in the marker interval AX-176791582_B02 - AX-176803063_B02, spanning 3.1 Mbp (**Fig. 2b**). The LOD score for MSF mapping was 53.3, and with the phenotypic variation explained (%PVE) of 61.3. A similar mapping was performed on the physical Tifruner genome sequence as a complementary analysis for the genetic mapping using the original 6871 SNPs (**Fig. 2C**). MSF was mapped to a locus at the end of Bo2 chromosome, in the marker interval AX-176805026_B02 - AX-176803063_B02, spanning 1.7 Mbp with –log10(p -Value= 59.5). These scores are suitable for the expected control of a major locus effect. BR was mapped to the same genomic location as MSF (**Fig. 2a**), with the same LOD score and %PVE, indicating that the same genetic factor controls both traits. BH was mapped on LG B05, in the marker interval AX-176823649_B05-AX-147223816_B05, spanning 2.2 Mbp (**Fig. 2a**), with a LOD score of 33.6, and %PVE of 46.6.

### Determination of MSF haplotype conservation in the US mini core collection

Two common SNP markers within the MSF region (118543259 to 120248910 bp) on LG B02 demonstrated polymorphism among accessions in the US mini core peanut collection. These markers separated the mini core accessions into two genotypic classes. Out of 112 accessions, 103 with high-quality data from the SNP array were included in this analysis. Forty-three accessions had the Hanoch SNP alleles, and 49 accessions had the IGC99 SNP alleles for the polymorphic loci. The remaining eleven accessions were excluded due to either recombination or heterozygosity/heterogeneity within this genomic region. The difference of MSF distribution between the Hanoch and IGHC99 alleles in the collection was inspected (**Fig. 3**). The distribution of MSF phenotype between the two allelic groups was highly significant (prob(ChiSq)<0.001), suggesting allelic conservation of the *MSF* locus in the collection. In the MSF-minus group, 32 accessions had the Hanoch allele, and only 4 had the IGC99 allele. In the MSF-plus group, 45 accessions had the IGC99 allele, and 11 had the Hanoch allele.

**Figure 3.**
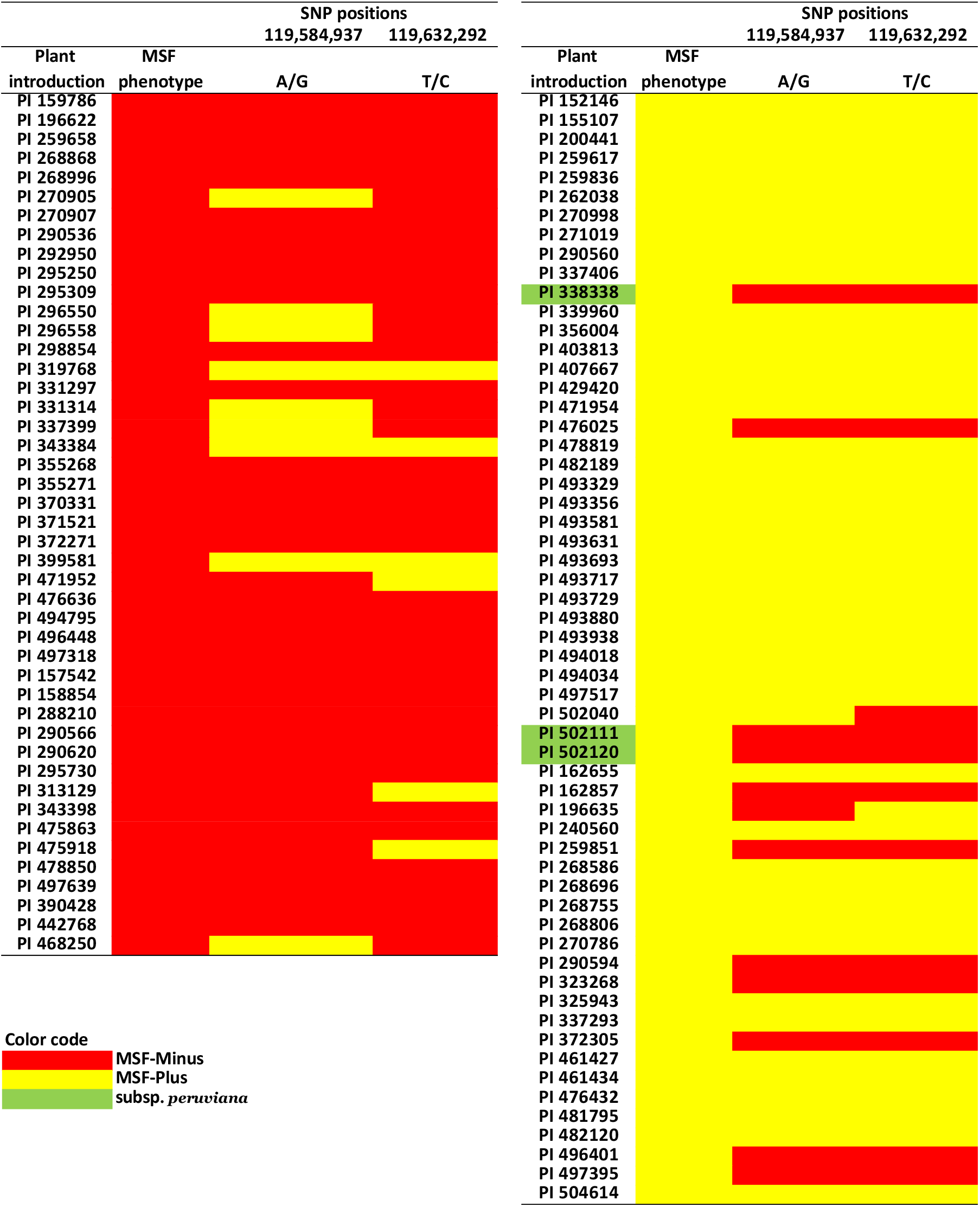
Conservative allele analysis of two SNP markers (AX-147227990, AX-176805840) across a mini-core collection of cultivated peanut. The collection was divided into two groups according to MSF phenotype: MSF-plus (yellow) and MSF-minus (red). If an allele of the genotype is in accordance with the MSF phenotype, it is marked with the same color. In green: three peruviana lines in the mini-core collection.

Interestingly, out of the 11 accessions with the MSF-plus phenotype and the Hanoch allele, 3 are accessions from *A. hypogaea* subsp. *peruviana* (green color in Fig. 3). Also, these are the only *peruviana* accessions in the mini core collection. Variety *peruviana* is considered an outgroup for the *hypogaea*/*fastigiata* cluster characterized by flowering on the main stem. So excluding this group from the analysis would further increase the significance of MSF distribution between the allelic groups.

### Identification of a TFL-1 like as a candidate gene for MSF

The mapping analyses resulted in a 1.7 Mbp fragment that encloses MSF. This region was subjected to a candidate gene analysis by extracting gene models from the tetraploid Tifrunner genome. A set of 168 genes was extracted from this genomic segment using the gene model order in PeanutBase (**Table S3**). Flowering locus protein T (IPR008914; Phosphatidylethanolamine-binding protein PEBP) (TFL1-like; CEN-like 1; Arahy.BBG51B) gene was found in the middle of this region (marked in yellow in Table S3). BLASTX of this gene with soybean (*Glycine max*) protein database showed that the best hit for *TFL-1* like (herein Ah*TFL1*) is Dt1 (81.94% identity), which previously was shown to control shoot determination (21).

The genomic DNA sequence of Ah*TFL1* protein was downloaded from the Tiffruner tetraploid genome and was compared with the tetraploid Shitouqi genome sequence (22), as well as with the diploid *Arachis durnensis* (A) and *Arachis ipaensis* (B) genomes (3) (**File S1**). Ah*TFL1* was found more similar to the A-genome homeologous sequence than the B-genome, although the trait was mapped to the B02 chromosome of the tetraploid genome. This is not surprising since the Ah*TFL1* is located in a middle of a region that was previously found to be AAAA tetrasomic region of cultivated peanut (19). Indeed, no difference was found between the A- and B-subgenome sequences of Ah*TFL1*. Furthermore, a 1492 bp deletion was found in the Shitouqi (*fastigiata*) genome sequence, spanning a 513 bp segment of the end of the Ah*TFL1* gene, leading to a truncated protein product (**Fig. 4a**). Interestingly, the deletion included a substantial part of the phosphatidylethanolamine-binding motif of the Ah*TFL1* gene (**Fig. 4a**).

**Figure 4.**
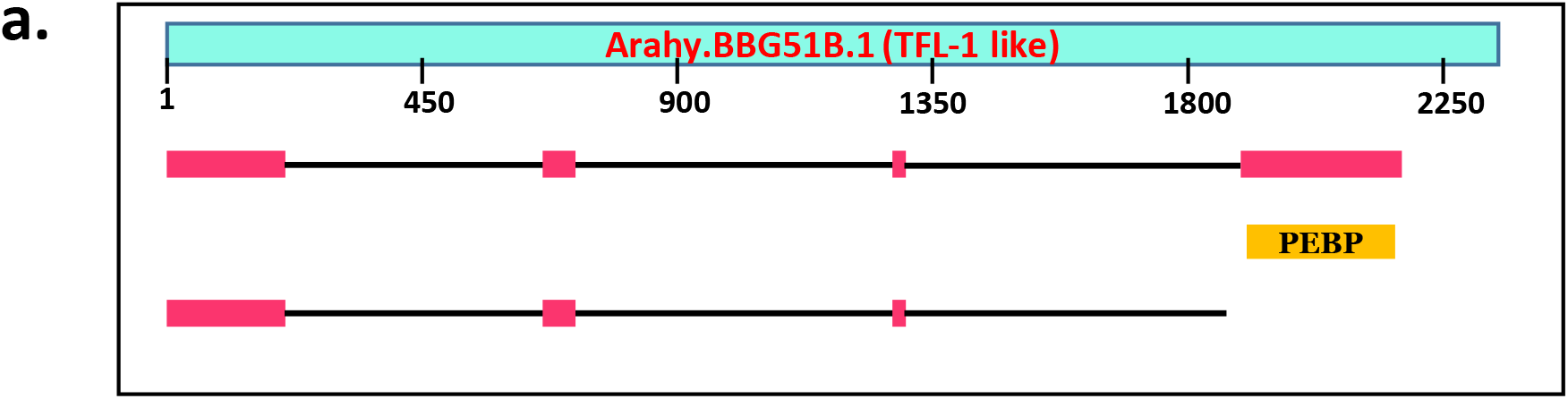
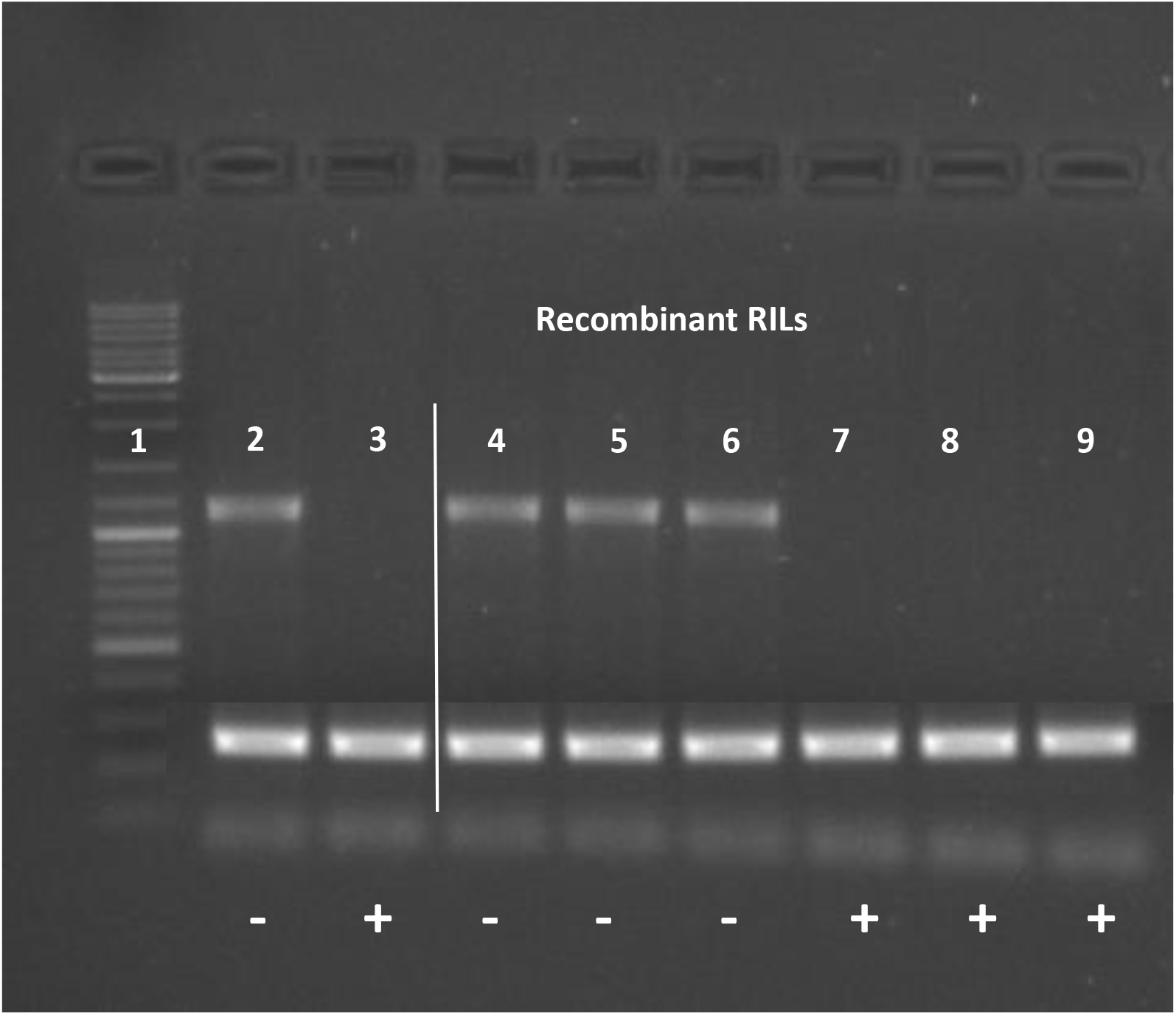
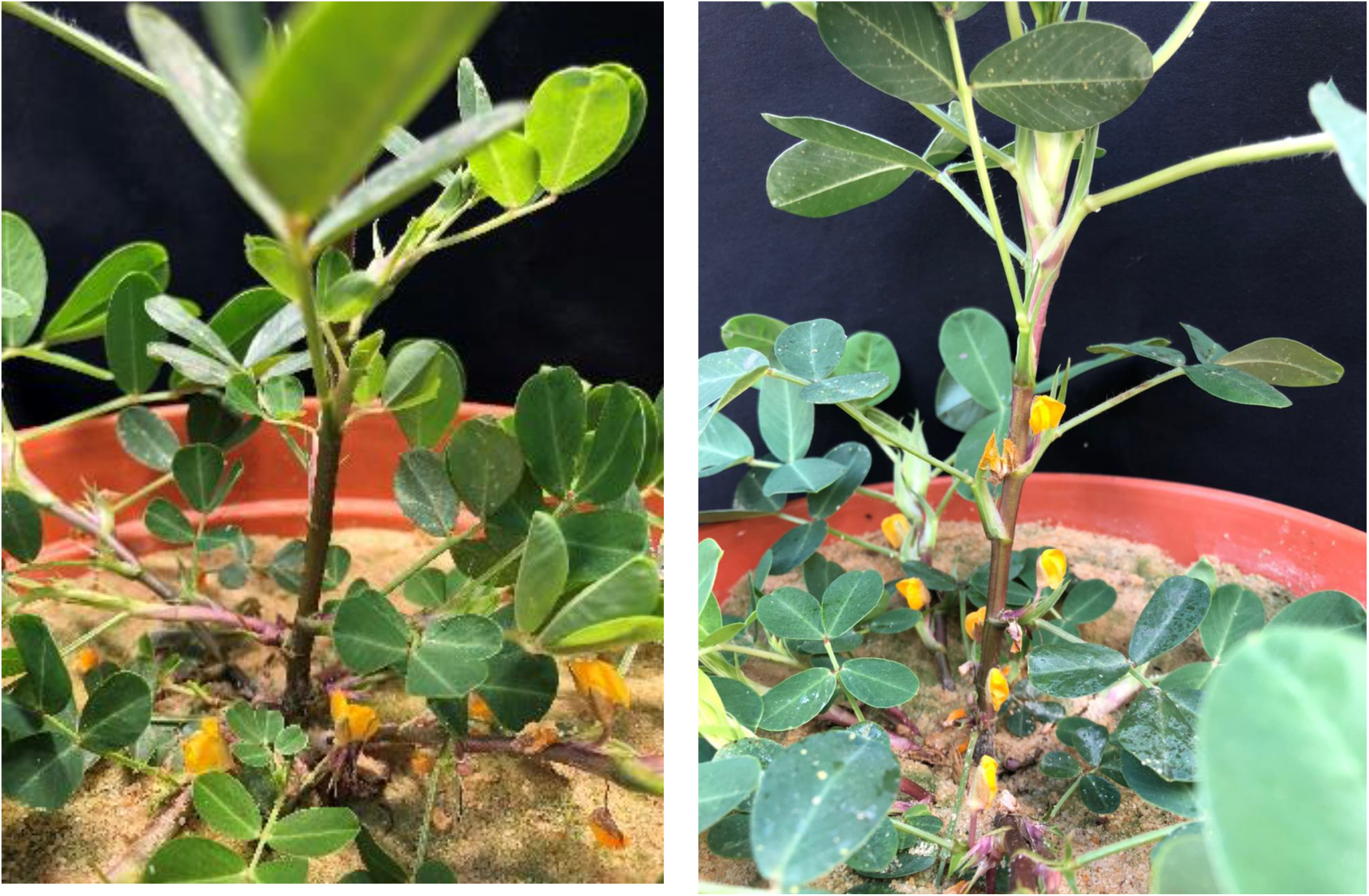
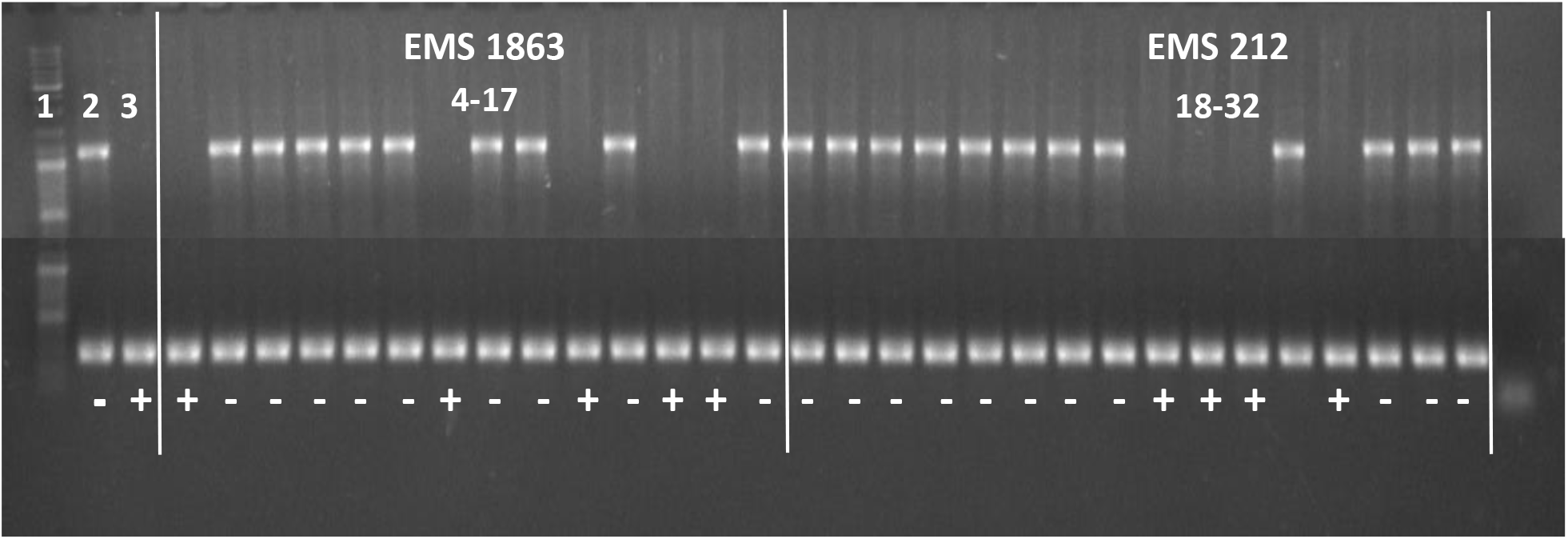
The deletion of *AhTFL1* in different peanut lines. **a**. Schematic illustration of the 513 bp deletion at the end of *AhTFL1* gene. Exons are indicated in pink; a black line indicates introns. **b**. Polymorphism of the 513 bp deletion of *AhTFL1* in Hanoch (lane 2), IGC99 (Lane 3), and RILs (4–9) that were recombinant for the original SNP markers and are not recombinant to the deletion. The figure shows 6 out of 37 recombinants in Hanoch X IGC99 RILs. + or − are indicating the MSF-plus and MST-minus phenotype. **c**. An example of two M2 individuals of the EMS 212 M2 family with MSF-plus and MSF-minus phenotype. d. . Polymorphism of the 513 bp deletion of *AhTFL1* in Hanoch (lane 2), IGC99 (Lane 3), and two segregating EMS M2 families (4–32). + or − are indicating the MSF-plus and MST-minus phenotype.

Based on the Ah*TFL1* genome sequence, specific primers were designed to capture the last exon of the Ah*TFL1* gene that includes the deletion. Polymorphism was found between the two parental lines Hanoch and IGC99, wherein the second had the deletion (**Fig. 4b**). In contrast to the sequence of Tifrunner, which had the complete Ah*TFL1* version in both A and B sub-genomes, Hanoch had one complete version and one truncated version of Ah*TFL1* (**Fig. 4b**). Remapping the MSF phenotype with the Ah*TFL1* deletion as a marker increased the LOD score from 53.3 to 158.8, and the –log10(p-Value) from 59.5 to 173.4, with no recombination in the RIL population (**Fig.4b**).

In order to acquire more evidence of Ah*TFL1* as the causative effect of MSF, we screened EMS M2 families that showed the MSF phenotype (**Fig. 4c**). Interestingly, two M2 families (EMS212 and EMS1863) that were segregating in 3:1 for MSF-minus:MSF-plus had the same Ah*TFL1* deletion (**Fig. 4d**). Also, MSF-plus plants did not develop secondary shoots. The deletion co-segregated with the MSF phenotype in each family (**Fig. 4d**).

To shed more light on the evolution of Ah*TFL1* in *Arachis*, in-silico sequence analysis was carried out. Re-sequencing genomic data of wild diploid, tetraploid landraces and tetraploid cultivars was retrieved from the SD2_Gerplasm_key of the Tifrunner genome sequencing publication (19). All diploids and tetraploid landraces did not have the complete deletion. In domesticated peanuts, out of 87 tested cultivars, in 79 (90.1%), the Ah*TFL1* deletion allele was according to the expectation. Still, several *A. hypogaea subsp*. *hypogaea* lines showed the deletion (data not shown).

## Discussion

In plants, the regulatory genes and proteins that control meristem determinacy and flowering are classified from the centrodialis/terminal flower/self-pruning (CETS) family, which are alternatively called phosphatidylethanolamine-binding proteins (PEBP) (23). In this family, *FLOWERING LOCUS T* (*FT*) and *TERMINALFLOWER I* (*TFL1*) of Arabidopsis and orthologs from other species are best-characterized controlling the balance and distribution of indeterminate and determinate growth. It has been suggested that FT promotes terminal flower development while TFL1 inhibits flowering (24, 25). FT also interacts with the basic leucine zipper transcription factor FD. FT and FD interaction can promote flowering, and overexpression of FD leads to branched sympodial growth (26). It was suggested that the domestication of many wild exotic plants into crops with desired growth habit might have resulted from the FT/TFL1 balances (23, 27–29). While this system has been investigated in legume crops (30, 31), it was never shown in peanut, in which branching pattern plays an important role in crop adaptability.

In this study, we have identified a *TFL1-like* gene that controls the flowering pattern in peanut. In wild and early domesticated peanuts, the alternative flowering pattern was the dominant phenotype. In this pattern, shoots that do not flower are generating secondary shoots, which are flowering. In such a biological system, it is expected that FT would promote flowering in all shoots, while TFL should be expressed in the tips of only non-flowering vegetative shoots. Indeed, searching the peanut expression Atlas (32, 33), the *AhTFL-1* was found to have an expression signature exclusively in non-flowering vegetative tips (**Fig. S2**). In the *fastigiata*-like “mutant”, *AhTFL-1* is truncated and therefore inactive. The FT/TFL ratio presumably is high, and the main shoots and the lateral shoots are forming flowers and not forming secondary shoots. Thus, the deletion in *AhTFL-1* in this study fits well into the central FT/TFL1 balance model.

Another question raised from the results is how *AhTFL-1* evolved in Arachis and how important it was for the formation of *fastigiata* types from *hypogaea* types. It was shown previously (Bertioli et al, 2019 (19), supplemental table 12-high-coverage-AB-SNP counts.xlsx deposited at https://doi.org/10.25739/hb5x-wx74) that an ancient tetrasomic event in all of the 39 cultivated peanut lines analyzed and *A. monticola*, the wild tetraploid relative of peanut, replaced a segment of chromosome B02 with A02 forming AA A’A’ genome composition from 118,258,164 to 120,571,497 bp . Therefore, this tetrasomic region encompasses *AhTFL-1* and preceded the deletion reported in this study. The deletion identified in this study occurred in one of the two sub-genomes (A or A’), similar to what we observed in in Hanoch. This change still would not lead to sequential flowering given the presence of functional alleles in one sub-genome. Finally, probably by another tetrasomic recombination event, the deletion spread to both AAA’A’, leading to sequential flowering, as it is presented in many cultivated *fastigiata* genotypes.

With regards to the importance of these changes in *AhTFL-1* for the historical event of subspecies *fastigiata*, we speculate that this gene had an important but not exclusive role. On the one hand, the significant haplotype conservation was found for the region adjacent to the *MSF* locus among two collections (USA mini core and the resequencing databases), suggesting an early selection upon *hypogaea*/*fastigiata* speciation. On the other hand, this conservation was not absolute, as several genotypes in both datasets were either MSF-plus without the deletion or MSF-minus with the presence of the deletion. Also, many of the *Arachis* landraces, i.e., *peruviana* and *hirsuta*, are MSF-plus but do not possess the deleted genotype of *AhTFL-1*. Therefore, several alternative pathways may lead to the MSF-plus phenotype in *Arachis*, and mutated *AhTFL-1* is not the exclusive one.

Another evidence for several possible pathways for MSF-plus phenotype comes from the EMS experiment. While two of the four inspected M2 families had the deletion, the other families did not. Moreover, in one family (EMS 995), the MSF-plus trait occurs already in the M1 generation, and all M2 individuals were MSF-plus, suggesting a dominant inheritance. A quick BLASTX analysis of the TFL/FT family in peanut revealed over 30 different copies of TFL-like genes in the tetraploid peanut genome (**Table S4**). Also, in a study that inspected the expression levels of 32 phosphatidyl ethanolamine-binding proteins in the wild diploid progenitors (34), *AhTFL1* was not one of the candidate genes with specific high expression levels in reproductive tips. In fact, small deletions in the *AhTFL1* coding sequence exist in the two wild progenitors (File S1). Therefore, other members of this family may have a role in the sequential flowering as well. The large TFL/FT family can also explain the relatively complicated branching pattern in peanut. As described above, the original phenotype of peanut is alternate flowering. There must be a mechanism that controls the flowering/non-flowering state of each alternative shoot in this pattern. Also, peanut’s shoot tips are indeterminate, regardless of the flowering pattern (i.e., MSF-plus or MSF-minus), in that a tip will continue to elongate and never terminate into a flower. This perennial nature of cultivated peanut was demonstrated in field-grown plants under protected conditions (35). Such a physiological system requires the involvement of a gene network, including subfunctionalization and neofunctionalization of genes from the TFL/FT family.

Interestingly, in this study, a large deletion was found in EMS-induced plants that perfectly matches the MSF phenotype. This is not an expected result, as EMS usually changes GC to AT and AT to GC. However, there is some evidence that EMS can cause base-pair insertions or deletions and more extensive intragenic deletions, especially at the end of chromosomes (36). Also, the *AhTFL-1* region is a hotspot for recombination events, evident by the large genetic distance but the relatively small physical distance between AX-147227990 and AX-176805840markers, with 37 recombinants in the RIL population. Therefore, the *AhTFL-1* region might be a hotspot for large mutational deletion that might be sensitive to chemical mutagenesis, which also played a role in peanut evolution. Alternatively, since a deletion of this size from EMS mutagenesis is unexpected, the result of the mutant lines could be a case of gene conversion process.

In conclusion, the integrated results from our study suggest that structure changes in the *AhTFL-1* gene have led to the formation of sequential flowering phenotype in peanut. Those events may also have played a historical role in the evolution of peanut speciation in domestication and modern cultivation.

---

**Figure S1.**
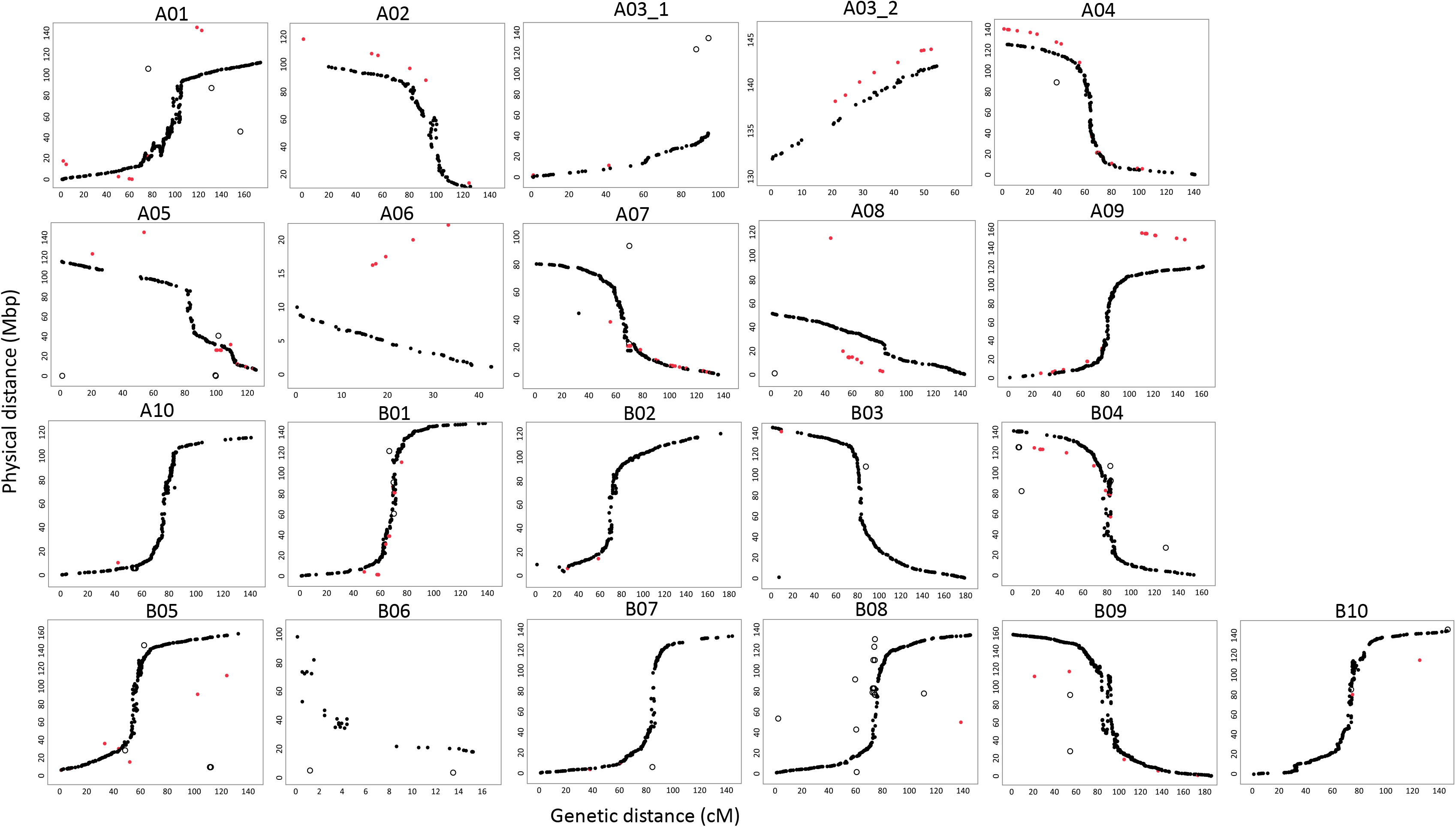
The correlation between the genetic distance (cM) (x-axis) of markers on each linkage group (LG) and the physical genome position (Mbp) (y-axis) based on the Tifrunner reference genome. Black dots represent the respective chromosome markers with the LG, red dots to the homologous chromosome.

**Figure S2.**
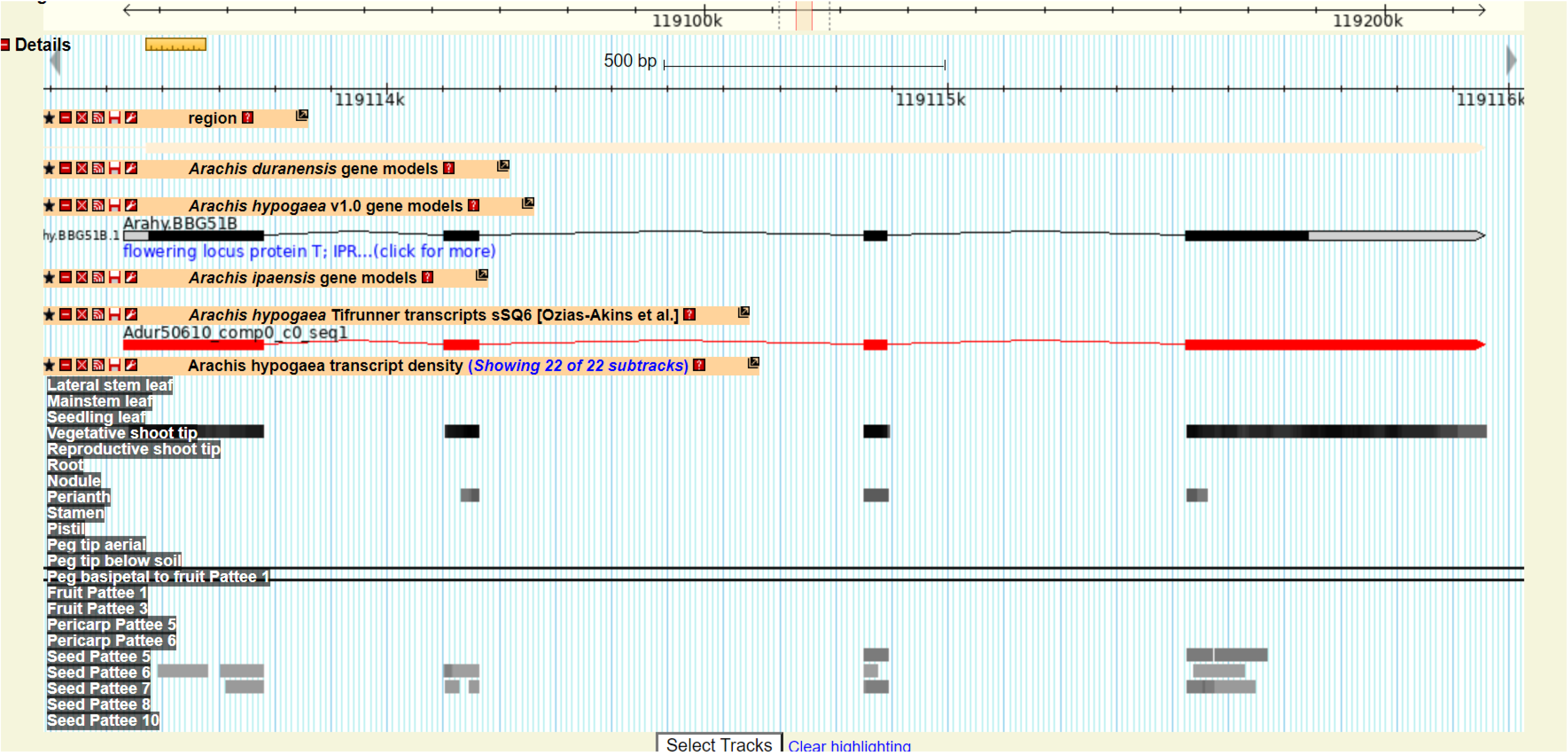
JBrows output of the *AhTFL1* gene, showing the Arachis hypogaea transcript density of 22 sub-tracks. Note for the almost exclusive expression of in the vegetative shoot tip.

**Table S1.** Phenotypic trait values of MSF, BR and BH in 269 Hanoch X IGC99 RILs.

**Table S2.** Hanoch X IGC99 RIL population genotyping, following the linkage map order, colored based on the alleles of the parents.

**Table S3.** A list of 168 gene models located at the MSF locus. AhTFL1 is marked in yellow.

**Table S4.** A list of TFL-like gene models in the tetraploid peanut genome.

**File S1**. Multiple alignment analysis to compare between the coding sequence of AhTFL1 protein in Tiffruner, Arachis durnensis (A), Arachis ipaensis (B) and Shitouqi (B) genomes.

## Cited Literature

1. FAOSTAT. Production quantities of Groundnuts, with shell by country 2019. Available from: http://www.fao.org/faostat/en/#data/QC/visualize.

2. Moretzsohn MC, Gouvea EG, Inglis PW, Leal-Bertioli SCM, Valls JFM, Bertioli DJ. A study of the relationships of cultivated peanut (*Arachis hypogaea*) and its most closely related wild species using intron sequences and microsatellite markers. Ann Bot-London. 2013;111(1):113–26.

3. Bertioli DJ, Cannon SB, Froenicke L, Huang GD, Farmer AD, Cannon EKS, et al. The genome sequences of *Arachis duranensis* and *Arachis ipaensis*, the diploid ancestors of cultivated peanut. Nat Genet. 2016;48(4):438–44.

4. Samoluk SS, Chalup L, Robledo G, Seijo JG. Genome sizes in diploid and allopolyploid *Arachis* L. species (section *Arachis*). Genet Resour Crop Ev. 2015;62(5):747–63.

5. Krapovickas A, Vanni RO. El maní de Llullaillaco. Bonplandia. 2009;18:51–5.

6. Krapovickas A, Gregory WC, Williams DE, Simpson CE. Taxonomy of the genus *Arachis* (Leguminosae). Bonplandia. 2007;16:7–205.

7. Chu Y, Chee P, Isleib TG, Holbrook CC, Ozias-Akins P. Major seed size QTL on chromosome A05 of peanut (*Arachis hypogaea*) is conserved in the US mini core germplasm collection. Mol Breeding. 2019;40(1):6.

8. Stalker HT, Simpson CE. Gerplasm resources in *Arachis*. In: Patee HE, Stalker HT, editors. Advances in Peanut Science. Stillwater, OK: American Peanut Research and Education Society; 1995. p. 14–55.

9. Pittman RN. United States peanut descriptors. US Department of Agriculture, Agricultural Research Service, ARS. 1995; 132.

10. Hammons RO. Inheritance of inflorescences in main stem leaf axils in *Arachis hypogaea* L. Crop Sci. 1971;11:570–1.

11. Jc W. Inheritance of branching pattern in *Arachis hypogaea* L. Peanut Sci. 1975(2):1–5.

12. Chopra R, Simpson CE, Hillhouse A, Payton P, Sharma J, Burow MD. SNP genotyping reveals major QTLs for plant architectural traits between A-genome peanut wild species. Mol Genet Genom. 2018;293(6):1477–91.

13. Hovav R, Hedvat I. (Hebrew) The development and screening of Hanoch-based EMS peanut population. A scientific report submitted to IGPB. 2011:1–11.

14. Clevenger JP, Korani W, Ozias-Akins P, Jackson S. Haplotype-based genotyping in polyploids. Front Plant Sci. 2018;9.

15. Korani W, Clevenger JP, Chu Y, Ozias-Akins P. Machine learning as an effective method for identifying true single nucleotide polymorphisms in polyploid plants. Plant Genome. 2019;12(1).

16. Patil A, Popovsky S, Levy Y, Chu Y, Clevenger J, Ozias-Akins P, et al. Genetic insight and mapping of the pod constriction trait in Virginia-Type peanut. Bmc Genet. 2018;19:93.

17. Van Ooijen JW, Voorrips RE. JoinMap version 4.1 software for the calculation of genetic linkage maps. In: Plant Research international B.V TN, editor. 2001.

18. Voorrips RE. MapChart 2.2: Software for the graphical presentation of linkage maps and QTLs. In: Plant Research International W, The Netherlands, editor. 2006.

19. Bertioli DJ, Jenkins J, Clevenger J, Dudchenko O, Gao DY, Seijo G, et al. The genome sequence of segmental allotetraploid peanut *Arachis hypogaea*. Nat Genet. 2019;51(5):877–84.

20. Van Ooijen JW. MapQTL 6. Software for the mapping of quantitative trait loci in experimental populations. In: Plant Research international B.V and Kyazma BV TN, editor. 2004.

21. Liu B, Watanabe S, Uchiyama T, Kong F, Kanazawa A, Xia Z, et al. The soybean stem growth habit gene *Dt1* Is an ortholog of arabidopsis *TERMINAL FLOWER1*. Plant Physiol. 2010;153(1):198–210.

22. Zhuang WJ, Chen H, Yang M, Wang JP, Pandey MK, Zhang C, et al. The genome of cultivated peanut provides insight into legume karyotypes, polyploid evolution and crop domestication. Nat Genet. 2019;51(5):865–76.

23. McGarry RC, Ayre BG. Manipulating plant architecture with members of the CETS gene family. Plant Sci. 2012;18:71–81.

24. Hanano S, Goto K. Arabidopsis TERMINAL FLOWER1 is involved in the regulation of flowering time and inflorescence development through transcriptional repression. Plant Cell 2011;23:3172–84.

25. Shalit A, Rozman A, Goldshmidt A, Alvarez JP, Bowman JL, Eshed Y, et al. The flowering hormone florigen functions as a general systemic regulator of growth and termination. Proc Nat Ac of Sci. 2009;106(20):8392–7.

26. Parmentier-Line CM, Coleman GD. Constitutive expression of the Poplar FD-like basic leucine zipper transcription factor alters growth and bud development. Plant Biotechnol J. 2015;14:260–70.

27. Kojima S, Takahashi Y, Kobayashi Y, Monna L, Sasaki T, Araki T, et al. Hd3a, a rice ortholog of the arabidopsis *FT* Gene, promotes transition to flowering downstream of *Hd1* under short-day conditions. Plant Cell Physiol. 2002;43(10):1096–105.

28. Blackman BK, Strasburg JL, Raduski AR, Michaels SD, Rieseberg LH. The role of recently derived *FT* paralogs in sunflower domestication. Curr Biol. 2010;20(7):629–35.

29. McGarry RC, Prewitt SF, Culpepper S, Eshed Y, Lifschitz E, Ayre BG. Monopodial and sympodial branching architecture in cotton is differentially regulated by the *Gossypium hirsutum SINGLE FLOWER TRUSS* and *SELF*◻*PRUNING* orthologs. New Pytol. 2016;212:244–58.

30. Tian Z, Wang X, Lee R, Li Y, Specht JE, Nelson RL, et al. Artificial selection for determinate growth habit in soybean. Proc Nat Ac Sci. 2010;107(19):8563–8.

31. Kwak M, Velasco D, Gepts P. Mapping homologous sequences for determinacy and photoperiod sensitivity in common bean (*Phaseolus vulgaris*). J Hered. 2008;99(3):283–91.

32. Dash S, Cannon EKS, Kalberer SR, Farmer AD, Cannon SB. PeanutBase and other bioinformatic resources for peanut. In: T. Sh, Wilson RF, editors. Peanuts Genetics, Processing, and Utilization. AOCS Press; 2016. p. 241–52.

33. Clevenger J, Chu Y, Scheffler B, Ozias-Akins P. A developmental transcriptome map for allotetraploid *Arachis hypogaea*. Front Plant Sci. 2016;7.

34. Jin H, Tang X, Xing M, Zhu H, Sui J, Cai C, et al. Molecular and transcriptional characterization of phosphatidyl ethanolamine-binding proteins in wild peanuts Arachis duranensis and Arachis ipaensis. BMC Plant Biol. 2019;19(1):484.

35. Kvien CK, Ozias-Akins P. Lack of monocarpic senescence in Florunner peanut. Peanut Sci. 1991;18:86–90.

36. Sega GA. A review of the genetic effects of ethyl methanesulfonate. Mutation Research/Reviews in Genetic Toxicology. 1984;134(2):113–42.

