## Supplementary material for "Identification of a Major Locus for Flowering Pattern Sheds Light on Plant Architecture Diversification in Cultivated Peanut": File S1

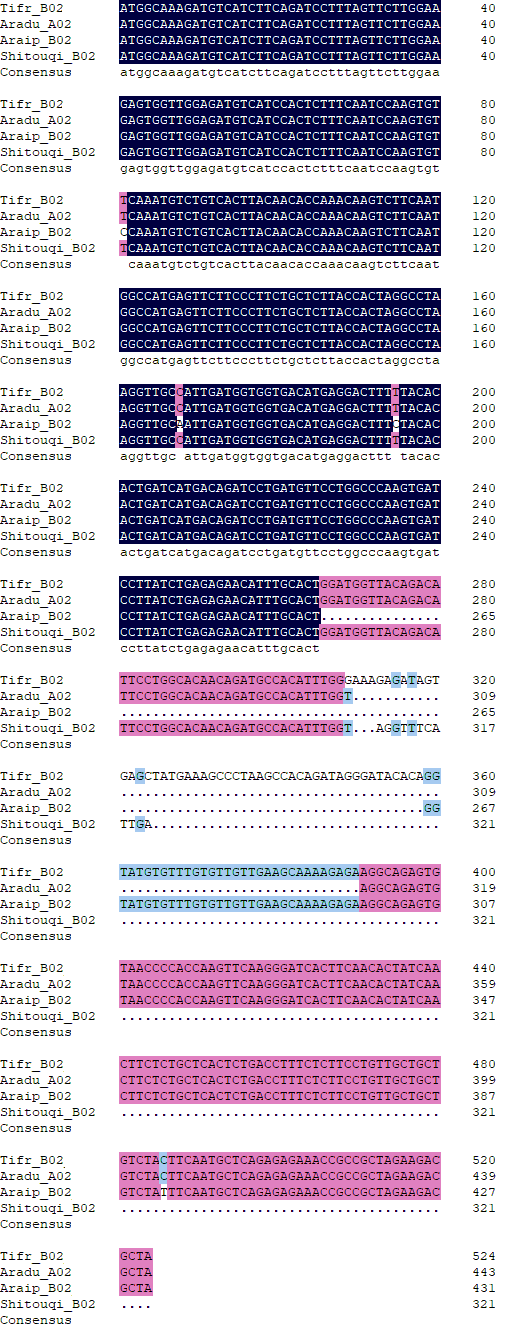

**File S1**. Multiple alignment analysis to compare between the coding sequence of Ah*TFL1* protein in Tiffruner, *Arachis durnensis* (A), *Arachis ipaensis* (B) and Shitouqi (B) genomes.
